# Rapid protocol for mitochondria isolation from cardiomyocytes employing cell strainer-based procedure

**DOI:** 10.64898/2026.04.02.716092

**Authors:** Joanna Lewandowska, Barbara Kalenik, Adam Szewczyk, Antoni Wrzosek

## Abstract

**Aims:** The development of a method for isolating mitochondria from a specific cell type within a given tissue, while preserving their structural and functional integrity to the greatest possible extent, remains an ongoing challenge. The aim of this study was to establish a protocol for the isolation of mitochondria from rodent cardiomyocytes, characterized by minimal contamination with other cell types and a high yield of mitochondrial fractions originating from distinct subcellular regions of cardiomyocytes.

**Methods and results:** In the present study, cardiomyocytes from guinea pig and rat hearts were isolated using a standard enzymatic digestion protocol in a Langendorff heart perfusion system. Traditionally, the isolation of organelles, including mitochondria, from whole cardiac tissue as well as from cardiomyocytes has relied primarily on mechanical tissue homogenization These conventional approaches involve the localized application of high pressure to cells, which may potentially damage delicate organelles, particularly mitochondria. Moreover, such homogenization preferentially releases mitochondria located in the subsarcolemmal region of cardiomyocytes rather than representing the entire mitochondrial population. In our study, we employed an alternative approach based on the gentle mechanical disruption of cardiomyocytes by passing the cell suspension through selected cell strainers using a cell scraper. This strategy facilitated mild disruption of cellular structures, significantly increasing the yield of mitochondria released from interfibrillar regions while preserving mitochondrial functionality. Moreover, this method decrease probability of sample contamination with mitochondria from other cells, based on cell size differences. The effectiveness of this method was confirmed by transmission electron microscopy, and high-resolution respirometry, which revealed no evidence of outer mitochondrial membrane damage, as indicated by the lack of response to the addition of exogenous cytochrome c to the incubation chamber. Moreover, mitochondrial oxygen consumption increased by 7.39 ± 1.25-fold following the addition of 100 µM ADP, reflecting efficient ADP-stimulated respiration. Furthermore, fluorescence measurements were performed. to assess changes in the mitochondrial inner membrane potential (ΔΨ). The isolated mitochondria were also suitable for electrophysiological studies using the single-channel patch-clamp technique. Additionally, mitochondria isolated using the protocol developed in our laboratory exhibited a high capacity for transplantation into H9c2 cells.

**Conclusion:** In summary, our mitochondrial isolation method is rapid, efficient, and yields functionally competent mitochondria. These preparations are suitable for a wide range of downstream applications, including patch-clamp electrophysiology, analyses of oxygen consumption under various pharmacological conditions, as well as mitochondrial transplantation.

**Graphical abstract:** 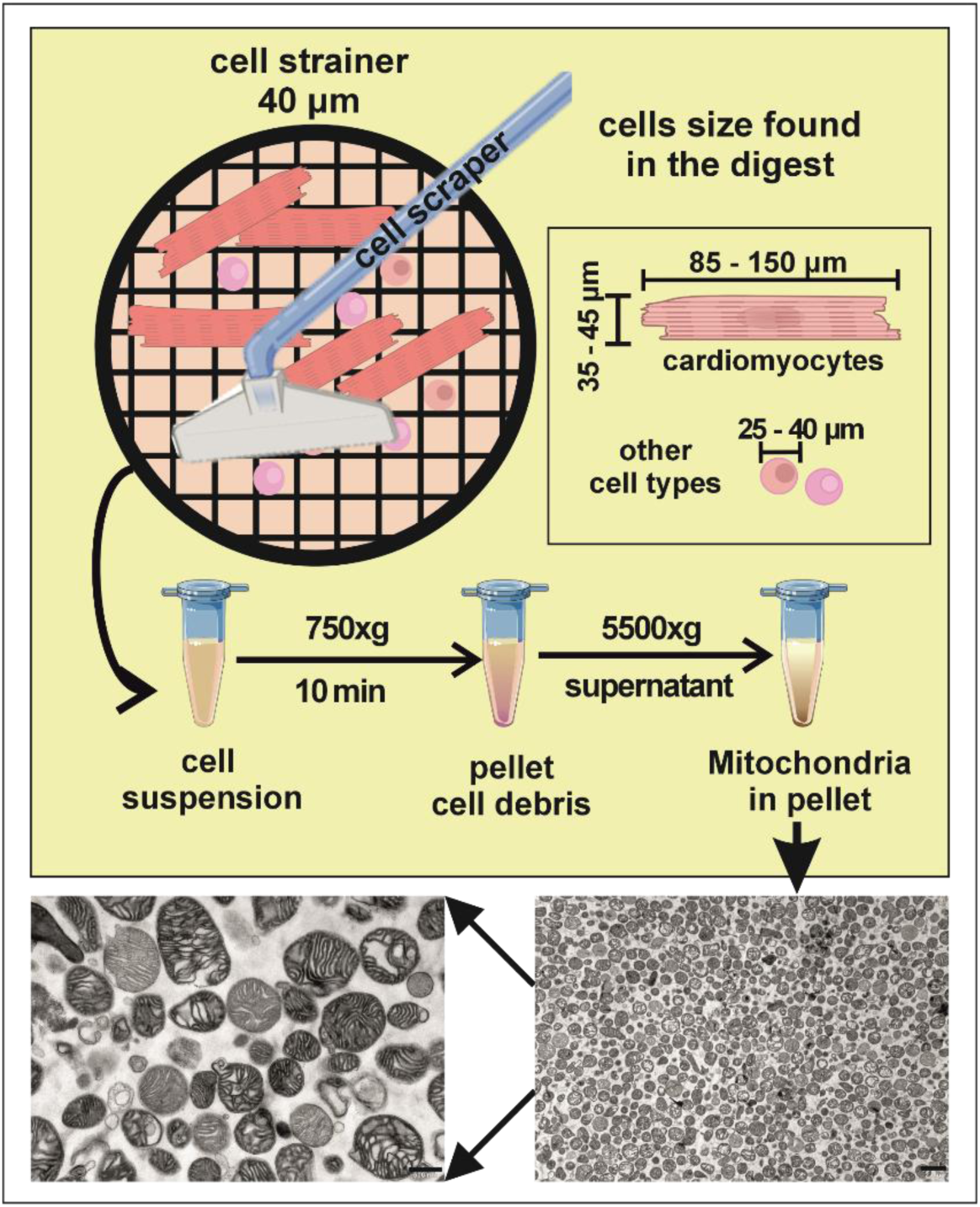

## 1. Introduction

History of mitochondria and isolated mitochondria research up to the 1980s are summarized in the review by Ernster and Schatz (Ernster & Schatz, 1981). This period includes discoveries assigning mitochondria the role of ATP-producing organelles through oxidative phosphorylation (Mitchell, 1961), as well as studies on the metabolism of normal and cancer cells (Warburg, 1956), (Spinelli & Haigis, 2018). Subsequent research demonstrated that mitochondria are not only organelles responsible for ATP synthesis, but also dynamic structures involved in cellular necrosis and apoptosis (Glover, Schreiner et al., 2024), as well as sites of reactive oxygen species (ROS) generation (Murphy, 2009), (Gómez, Mota-Martorell et al., 2023), (Hernansanz-Agustín & Enríquez, 2021). Mitochondrial ROS play dual roles, functioning in redox signaling and in oxidative stress, during physiological and pathological conditions, respectively. Later new mitochondrial functions have been discovered, including regulation of intracellular Ca²⁺ levels (Mammucari, Raffaello et al., 2018), (Modesti, Danese et al., 2021), gene expression, metabolic adaptation, and stress response (Quirós, Mottis et al., 2016), (Brand, 2016), (Chandel, 2015). It has also been observed that mitochondria can be transferred between cells (Spees, Olson et al., 2006), (Liu, Mao et al., 2024). Another significant aspect is the possibility of transplanting isolated mitochondria as an innovative form of cellular biotherapy (Bian et al., 2026), (Lu et al., 2025), (Ali Pour, Hosseinian et al., 2021), (Ali Pour, Kenney et al., 2020), especially in primary and secondary mitochondrial diseases (Niyazov, Kahler et al., 2016), (Valenti & Vacca, 2022). It appears that most discoveries are based on preparations of isolated mitochondria, both in early studies (Claude, 1946), (Ernster & Schatz, 1981), and in contemporary research (Alia et al., 2026), (Bian et al., 2026), (Liu et al, 2026), (Lu et al., 2025). Obtaining functional mitochondria from cells and tissues is crucial for preserving their physiological state. Currently, numerous publications address this topic (Huang, Jin et al., 2024); (Djafarzadeh & Jakob, 2017); (Liao, Bergamini et al., 2020), and companies such as Luca Science LTD are attempting to patent organelle isolation methods. Based on the available literature, key steps in mitochondrial isolation include homogenization methods, appropriate buffer selection and temperature control (Kumari & Sadeesh, 2024). Glass homogenizers with varying clearance, adapted to specific cell and tissue types, are most commonly used for homogenizing isolated cells. Dounce (Dounce et al., 1955), Potter homogenizers, including the Braun S homogenizer (Djafarzadeh & Jakob, 2017), (Liao et al., 2020), as well as commercial meat grinders and Waring blenders are utilized. Unfortunately, homogenization may damage mitochondria (Brustovetsky, Jemmerson et al., 2002), (Gottlieb & Adachi, 2000). Differential centrifugation constitutes another essential step in most mitochondrial isolation methods. In some cases, detergents have also been used. One example is digitonin, applied for mitochondrial isolation (Dixit, Vanhoozer et al., 2021); however, it may disrupt mitochondrial membranes. Therefore, some authors recommend its use primarily for isolating mitochondrial subfractions, such as the outer mitochondrial membrane, mitoplasts, or mitochondrial DNA (Huang et al., 2024), (Howell, Nalty et al., 1986), (Nishimura & Yano, 2014). It was shown that low concentrations of digitonin weaken the plasma membrane, enabling cell disruption and mitochondrial isolation through gentle pipetting and differential centrifugation. Although this method did not appear to damage mitochondria, its efficiency was lower than that of traditional homogenization methods. Additional purification steps have also been proposed, such as density gradient centrifugation using Percoll (Anunciado-Koza, Guntur et al., 2023), (Wieckowski, Giorgi et al., 2009). Furthermore, purification using magnetic beads conjugated with anti-TOMM22 and anti-TOMM20 antibodies has been introduced (Hornig-Do, Günther et al., 2009), (Huang et al., 2024).

As mentioned above, homogenization is usually a critical step in obtaining functional mitochondria. Unfortunately, this method causes intensive shear-stress what can influence mitochondrial functions.

In the present study, we describe a method for isolating mitochondria from adult guinea pig and rat cardiomyocytes. The method is based on passing isolated cardiomyocytes through a cell strainer. We show, that it provides mitochondria suitable for a wide range of downstream applications, like analyses of oxygen consumption under various pharmacological conditions, patch-clamp electrophysiology, as well as mitochondrial transplantation. In summary, our mitochondrial isolation method is rapid, efficient, and yields fully functional mitochondria.

## 2. Methods

The procedure for isolating mitochondria from cardiomyocytes involves enzymatic digestion of the heart using the Langendorff perfusion method. The obtained cardiomyocytes are then gently disrupted on a cell strainer with a 40 μm diameter opening. A schematic diagram of the mitochondrial isolation process and its key components is presented in Figure 1.

**Figure 1.**
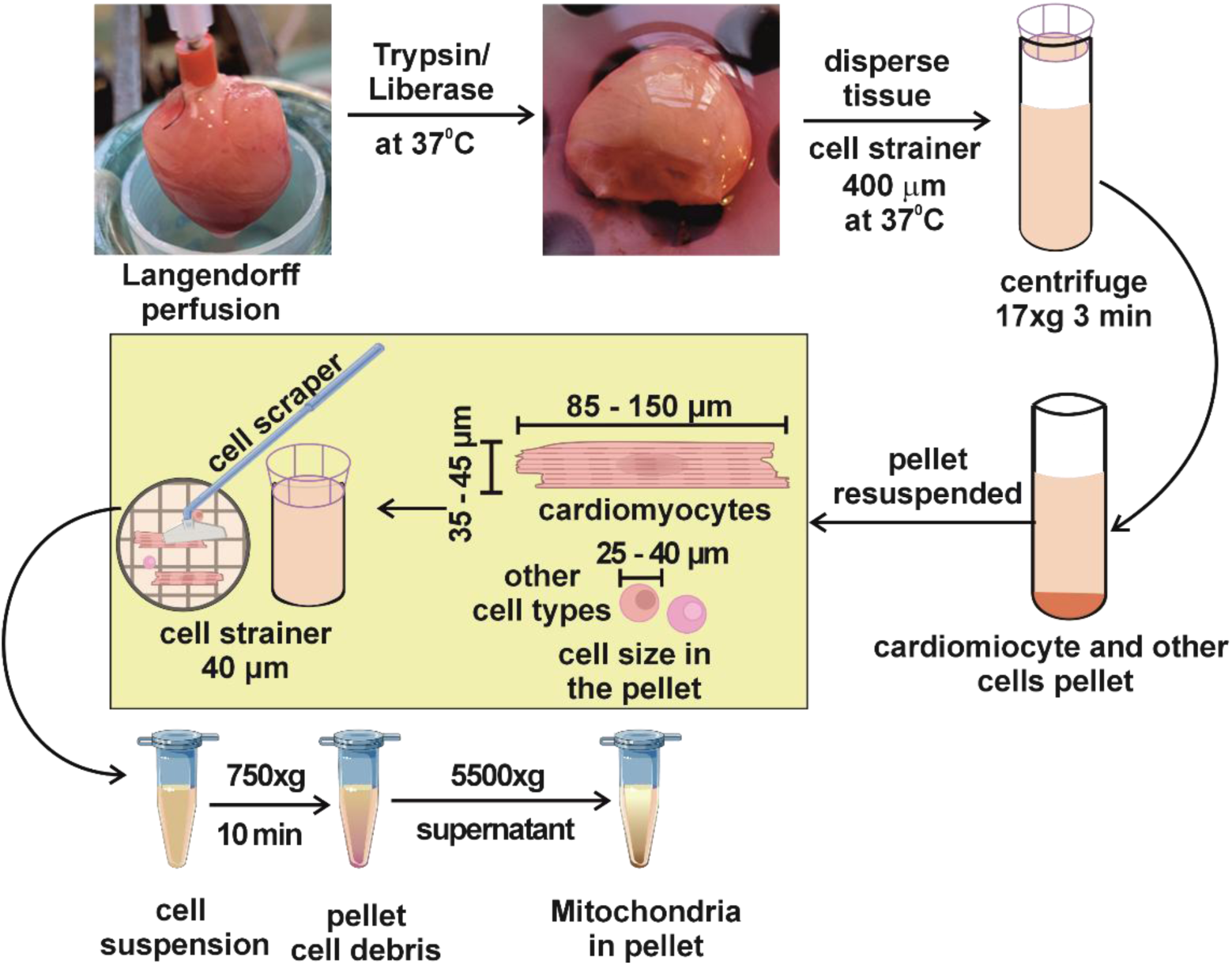
Experimental approach to isolate mitochondrial fraction from rodent cardiomyocytes. Schematic representation of a procedure for mitochondria isolation. Cardiomyocytes were liberated with digestion on Langendorff perfusion system. Larger fragments of connective tissue were removed by filtration through a cell strainer with a pore size of 400 µm and cells were pelleted by centrifugation. Than cardiomyocytes were gently disrupted using a cell scraper on a cell strainer with a pore size of 40 µm and cardiomyocytes mitochondria together with other cardiac cells were passing the cell suspension through strainer pore. Then cell suspension is centrifugated at 750xg and cardiomyocytes mitochondria remain suspended, while other cardiac cells are located in discarded pellet. At the final step, cardiomyocytes mitochondria are pelleted and resuspended in a buffer and concentration needed for further studies.

### 2.1 Animals

All investigations involving animals conformed to the Guidelines for the Care and Use of Laboratory Animals published by the US National Institutes of Health, and the experimental procedures used in this study were approved by the local Animal Research Committee of the Nencki Institute of Experimental Biology.

Guinea pig Dunkin Hartley - SPF HsdDhl:DH were purchased from AnLab s.r.o. (Czech Republic) weighing below 200g and grown later in our animal house. Males weighing 350g - 650g were used for experiments.

### 2.1 Cardiomyocytes Isolation

Cardiomyocytes were isolated as previously described (Frankenreiter, Bednarczyk et al., 2017), with minor modifications. Briefly, the heart excised from the animal was immediately placed in a perfusion buffer (composition: 113 mM NaCl, 25 mM KCl, 0.1 mM KH_2_PO_4_, 0.6 mM Na_2_HPO_4_, 1.2 mM MgSO_4_, 0.03 mM Phenol red, 11.6 mM NaHCO_3_, 10.1 mM KHCO_3_, 10 mM HEPES, 30 mM Taurine, 500 mM 2,3-Butanedione monoxime, 1 g/l glucose, pH 7.4) prewarmed to 37°C and gently compressed for approximately 5 seconds to remove excess blood, after which it was transferred to buffer maintained at 4°C. Under a stereomicroscope, the prepared heart was cannulated via the aorta above the aortic valve, and secured to the cannula using 5-0 silk sewing thread. Perfusion was initiated with described above perfusion buffer at a flow rate of 2.5 mL/min for approximately 10 minutes to remove residual blood from the coronary circulation. Subsequently, the buffer was replaced with perfusion buffer containing digestive enzymes, namely liberase and trypsin. Next similar buffer with trypsin and DNase was used. Enzymatic digestion was continued until a visible change in cardiac tissue structure was observed (Figure 1), typically after approximately 10 minutes. The heart was then removed from the cannula, and at 37°C the atria and remaining portions of the aorta were excised. The ventricles were cutted into small fragments using scissors and gently triturated with a wide-bore Pasteur pipette. Larger fragments of connective tissue were removed by filtration through a cell strainer with a pore size of 400 µm, then digestion was terminated by the addition of 2.5 mL perfusion buffer supplemented with 10% fetal bovine serum (FBS) per 30 mL of cell suspension. The resulting cardiomyocytes were pelleted by centrifugation at 19 × g for 3 minutes at room temperature. The obtained cell pellet was then resuspended in buffer appropriate for mitochondrial isolation.

### 2.2 Mitochondrial Isolation

Mitochondria were isolated from cardiomyocytes in a buffer composed of 250 mM sucrose, 5 mM HEPES, and 1 mM EGTA (pH 7.2), supplemented with 1% bovine serum albumin (BSA). Cells were mechanically disrupted by gentle trituration in portions through a 40 µm mesh filter continuously immersed in homogenization buffer. The resulting filtrates were centrifuged at 750 × g for 10 minutes.

To increase yield, the pellet was resuspended in homogenization buffer and passed through a 30 µm mesh filter. The obtained filtrate was again centrifuged at 750 × g for 10 minutes. The combined supernatants were subsequently centrifuged at 5,500 × g for 6 minutes. The final pellet, exhibiting a light brown coloration, contained mitochondria and was used for further analyses. All procedures were performed on ice, and all centrifugation steps were carried out at 4°C.

### 2.3 Electron microscopy

All samples for TEM imaging were prepared according to the same protocol. If not indicated otherwise, each step was followed by centrifugation 2 min, 6000 rpm. Briefly, pellets were washed 6x with 0.1M cacodylate buffer. Then, samples were treated with 2% osmium tetroxide in 0.1M cacodylate buffer, 60 min in the darkness. After 3 x washing with water samples were treated with 2% uranyl acetate in water, 40 min in the darkness. After 3 x washing with water samples were dehydrated for 10 min in each concentration of acetone dilutions: 25%; 50%; 75%; 90%; 96% and finally 15 min in 100% acetone. Then samples were impregnated by resin - acetone:epon resin 1:1, 1h and acetone:epon resin 1:2, 1h; epon resin 1h; epon resin o/n. During steps of resin impregnation centrifugation 2 min, 7000 rpm was used. Then pellets were polymerased in 60°C for 3 days. Next, samples were cutted on ultramicrotome (70 nm sections), contrasted with 2% lead citrate and viewed under a microscope.

Samples for TEM imaging and electron microscopy images were performed by the Laboratory of Electron Microscopy (Nencki Institute of Experimental Biology PAS) with high-resolution transmission electron microscope JEM 1400 (JEOL Co., Japan, 2008) with 11 Megapixel TEM Morada G2 camera (EMSIS GmbH, Niemcy).

### 2.4 Cell culture conditions

The rat embryonic cardiomyoblast-derived cell line H9c2 was obtained from ATCC. The cells were cultured at 37°C in a humidified atmosphere containing 5% CO_2_ in DMEM supplemented with 10% fetal bovine serum, 4 mM glutamine, 100 U/ml penicillin, and 100 μg/ml streptomycin.

### 2.5. Visualization of the cell mitochondria and measurements of inner mitochondrial membrane potential

Cardiomyocytes and H9c2 cells were loaded with the fluorescent dyes MitoTracker™ Green FM, MitoRed, or JC-1 (Invitrogen) according to the manufacturer’s protocols in culture medium at 37°C in a humidified atmosphere containing 5% CO_2_ for 20 minutes. Subsequently, the incubation medium was replaced with FluoroBrite™ DMEM (Gibco). Fluorescence imaging was performed using an Olympus IX83 confocal microscope equipped with a 60× oil-immersion objective (UPLSAPO 60XS). Image analysis was carried out using ImageJ software.

### 2.6 Mitochondrial membrane potential measurements

The measurements were made at room temperature in the 1 mL cuvette of a FluoroMax-4 spectrofluorometer (Horiba Scientific) using 1 μM Rhodamine 123, a membrane potential sensitive fluorescence dye. The measurements were performed in a medium Miro 5 containing 10 mM KH_2_PO_4_, 60 mM KCl, 60 mM Tris, 5 mM MgCl_2_, 110 mannitol and 0.5 mM EDTA, pH 7.4. As respiratory substrates 10 mM glutamate, 5 mM malate, and 5 mM succinate were used. The samples were excited at 450 nm and the fluorescence was registered at 550 nm.

### 2.7 Mitochondrial respiration

Mitochondrial oxygen consumption was measured at 37°C using an Oroboros oxygraph (Austria) in a above described Miro 5 medium. As respiratory substrates 10 mM glutamate, 5 mM malate, and 5 mM succinate were used.

### 2.8 Patch-clamp experiments

Patch-clamp experiments were performed as previously described (Kampa et al., 2021; Maliszewska-Olejniczak et al., 2024; Walewska et al., 2022). Briefly, mitochondria isolated from guinea pig cardiomyocytes were converted into mitoplasts by incubation in a hypotonic solution containing 5 mM HEPES and 100 μM CaCl_2_, pH 7.2 for approximately 2 minutes to induce swelling and rupture of the outer mitochondrial membrane. Subsequently, a hypertonic solution composed of 750 mM KCl, 30 mM HEPES and 100 μM CaCl_2_, pH 7.2 was added to restore isotonic conditions. The final isotonic bath solution contained 150 mM KCl, 10 mM HEPES, and 100 μM CaCl_2_ pH 7.2. Patch-clamp pipettes were also filled with the same isotonic solution. Mitoplasts were readily identified based on their size, rounded morphology, transparency, and the presence of a characteristic “cap,” which distinguished them from other residual structures present in the preparation. In all control conditions, the isotonic solution contained 100 μM CaCl_2_. A low-calcium isotonic solution used for mitoBK_Ca_ channel characterization contained 150 mM KCl, 10 mM HEPES, 1 mM EGTA, and 0.752 mM CaCl_2_ (yielding 1 μM free Ca^2+^, pH 7.2). Channel modulators were applied as dilutions in isotonic solutions containing either 100 μM or 1 μM Ca^2+^. Compound delivery was achieved using a perfusion system consisting of a holder equipped with a custom-made glass capillary, a peristaltic pump, and Teflon tubing. Mitoplasts attached to the tip of the recording pipette were transferred into the wells of a multi-barrel perfusion system, where their external surfaces were continuously perfused with experimental solutions (Figure 6A). All experiments were conducted in the inside-out patch-clamp configuration. Reported voltages correspond to potentials applied to the interior of the patch-clamp pipette, with positive potentials reflecting the physiological polarization of the inner mitochondrial membrane (positive potential on the outside). Electrical connections were established using Ag/AgCl electrodes, with agar bridges (3 M KCl) serving as reference electrodes. Currents were recorded using a patch-clamp amplifier (Axopatch 200B, Molecular Devices, USA). Borosilicate glass pipettes with resistances of 10–20 MΩ were fabricated using a Narishige PC-10 puller. Signals were low-pass filtered at 1 kHz and sampled at 100 kHz.

### 2.9 Transplantation of mitochondria

Mitochondria in H9c2 cells and rat cardiomyocytes were labelled with MitoTracker Green and MitoRed dye, respectively, in conditions described in chapter 2.5 for 30 min. Then mitochondria were isolated with the method described in this article. Next, mitochondrial suspension was added to H9c2 cells cultured in 35mm dishes. After centrifugation (200 x g, 10 min, RT) cells were grown for about 16 h. Then labeled mitochondria were visualized with method described in 2.5 chapter.

## Results

If we isolated mitochondria from 1 mln of cell we obtained on average 11.3 mg of mitochondrial proteins.

### Structural Integrity and Mitochondrial Organization in Isolated Cardiomyocytes

Cardiomyocytes isolated using the Langendorff perfusion method were suspended in culture buffer and stained with MitoTracker™ Green FM to assess mitochondrial organization (Figure 2A). The dye accumulated within mitochondria in a manner dependent on the inner mitochondrial membrane (IMM) transmembrane potential (ΔΨ), revealing a highly ordered intracellular pattern characteristic of functionally intact cardiomyocytes. The spatial distribution of fluorescence exhibited a periodicity of 1.81 ± 0.06 μm, consistent with the canonical arrangement of mitochondria between myofibrils. Light microscopy demonstrated that the majority of isolated cardiomyocytes retained a rod-shaped morphology, indicative of preserved structural integrity and absence of hypercontracture (Figure 2B1). Transmission electron microscopy (TEM) confirmed preservation of mitochondrial ultrastructure, including well-defined cristae, and demonstrated their regular alignment with the contractile apparatus (Figure 2B2). Mitochondrial fraction obtained with our isolation protocol consisted predominantly of intact mitochondria (Figure 2B3). High-magnification TEM revealed well-preserved internal architecture comparable to that observed in situ (Figure 2B4).

**Figure 2.**
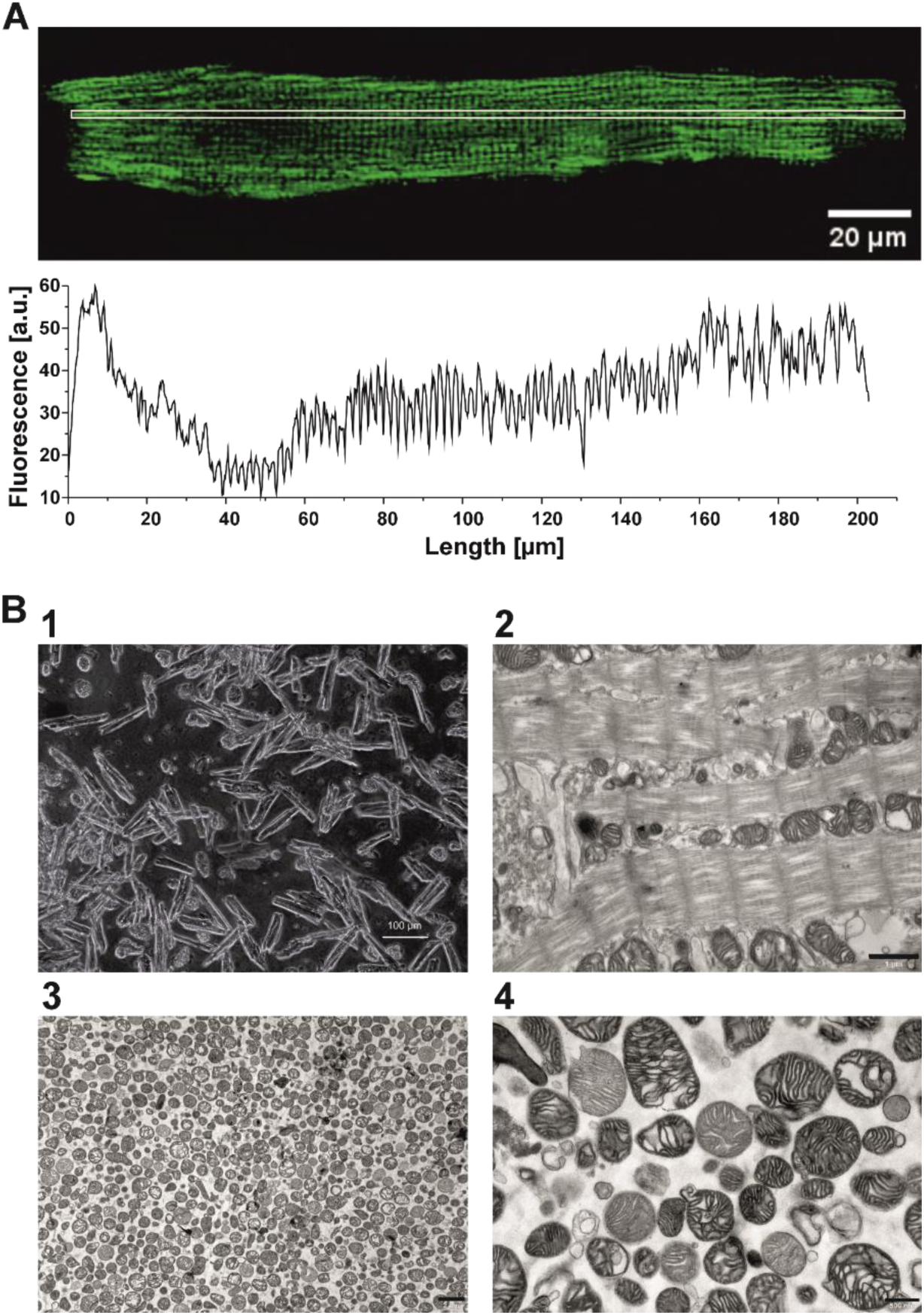
Microscopic analysis of isolated cardiomyocytes and mitochondrial fraction from guinea pig cardiomyocytes. (A) Fluorescence of cardiomyocyte labeled with MitoTracker Green; (B) light microscopy of isolated cardiomyocytes [1], electron microscopy of isolated cardiomyocytes [2] and isolated mitochondria [3, 4]. Scale bar 100 um; 1um; 2um and 500nm for [1]; [2]; [3]; and [4], respectively.

**Figure 3.**
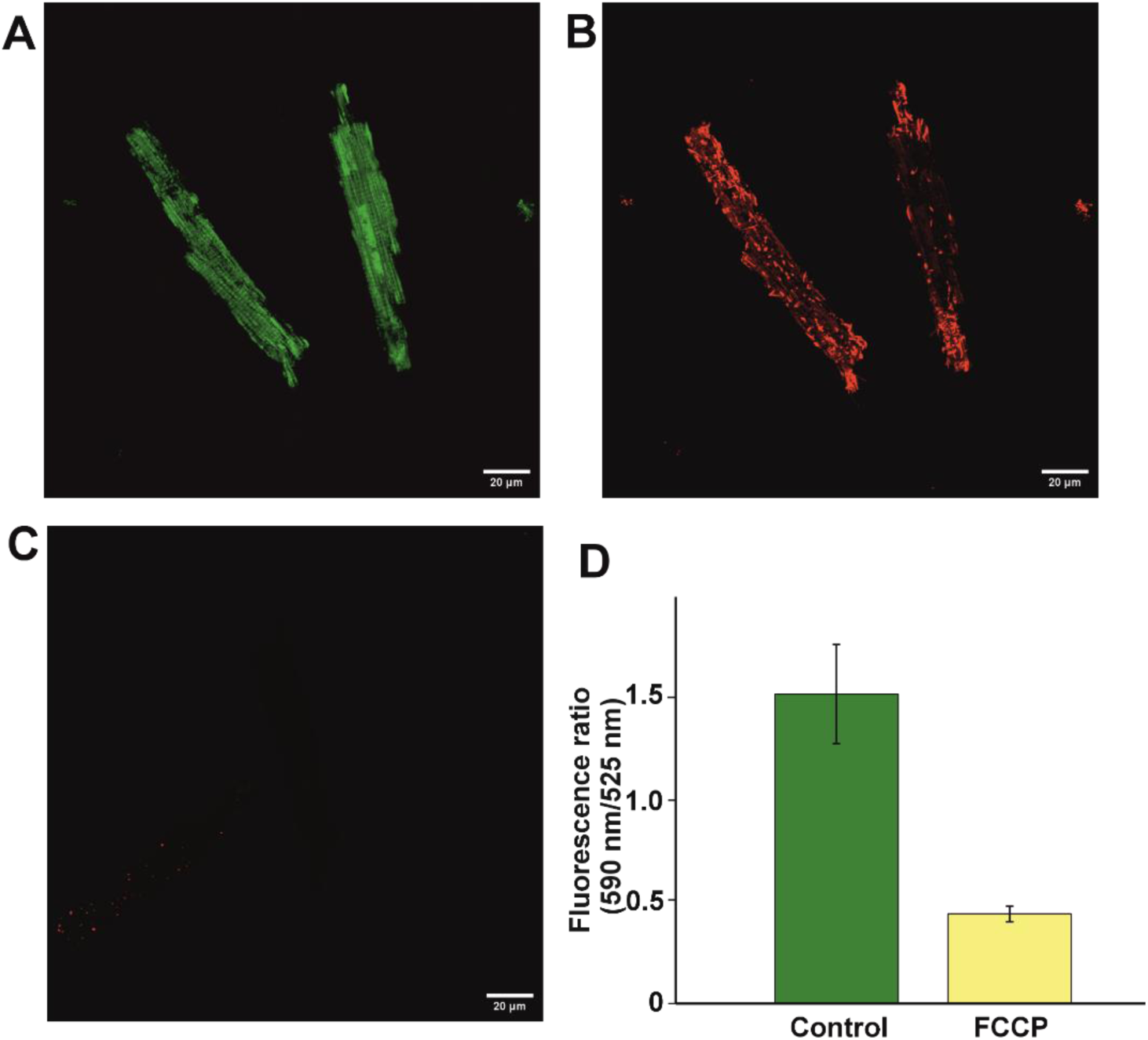
Measurement of the inner mitochondrial membrane potential using JC1. (A) Monomer fluorescence, (B) J-aggregate fluorescence, (C) Inner mitochondrial membrane potential in the presence of FCCP, (D) J-aggregate/monomer fluorescence ratio.

### Preservation of Mitochondrial Membrane Potential in Isolated Cardiomyocytes

Mitochondrial function was evaluated using the mitochondrial membrane potential probe JC-1. Under basal conditions, mitochondria exhibited predominant red fluorescence corresponding to J-aggregate formation, indicative of a high transmembrane potential ΔΨ (Figure 2A,B). Application of the protonophore FCCP (1 μM) resulted in a complete loss of J-aggregate fluorescence and a shift toward the monomeric green signal, reflecting mitochondrial depolarization (Figure 2C). Quantitative analysis of the red-to-green fluorescence ratio confirmed a marked dissipation of ΔΨ in the presence of FCCP (Figure 2D).

### Functional Characterization of Isolated Mitochondria by ΔΨ Measurements

The functional integrity of isolated mitochondria was assessed fluorometrically using Rhodamine 123 dye in MiR05 buffer supplemented with respiratory substrates (malate/glutamate and succinate). Addition of mitochondria resulted in rapid quenching of fluorescence, consistent with ΔΨ-dependent dye accumulation into mitochondria and self-quenching. Subsequent addition of ADP (150 μM) induced a transient increase in fluorescence, reflecting partial depolarization of ΔΨ during state III respiration, followed by recovery upon ADP phosphorylation to ATP (state IV). Application of FCCP (1 μM) caused a sustained increase in fluorescence, consistent with complete dissipation of ΔΨ. Representative recordings and pooled data are presented in Figure 4A and 4B, respectively. These ΔΨ dynamics indicate preserved functionality of the electron transport chain.

**Figure 4.**
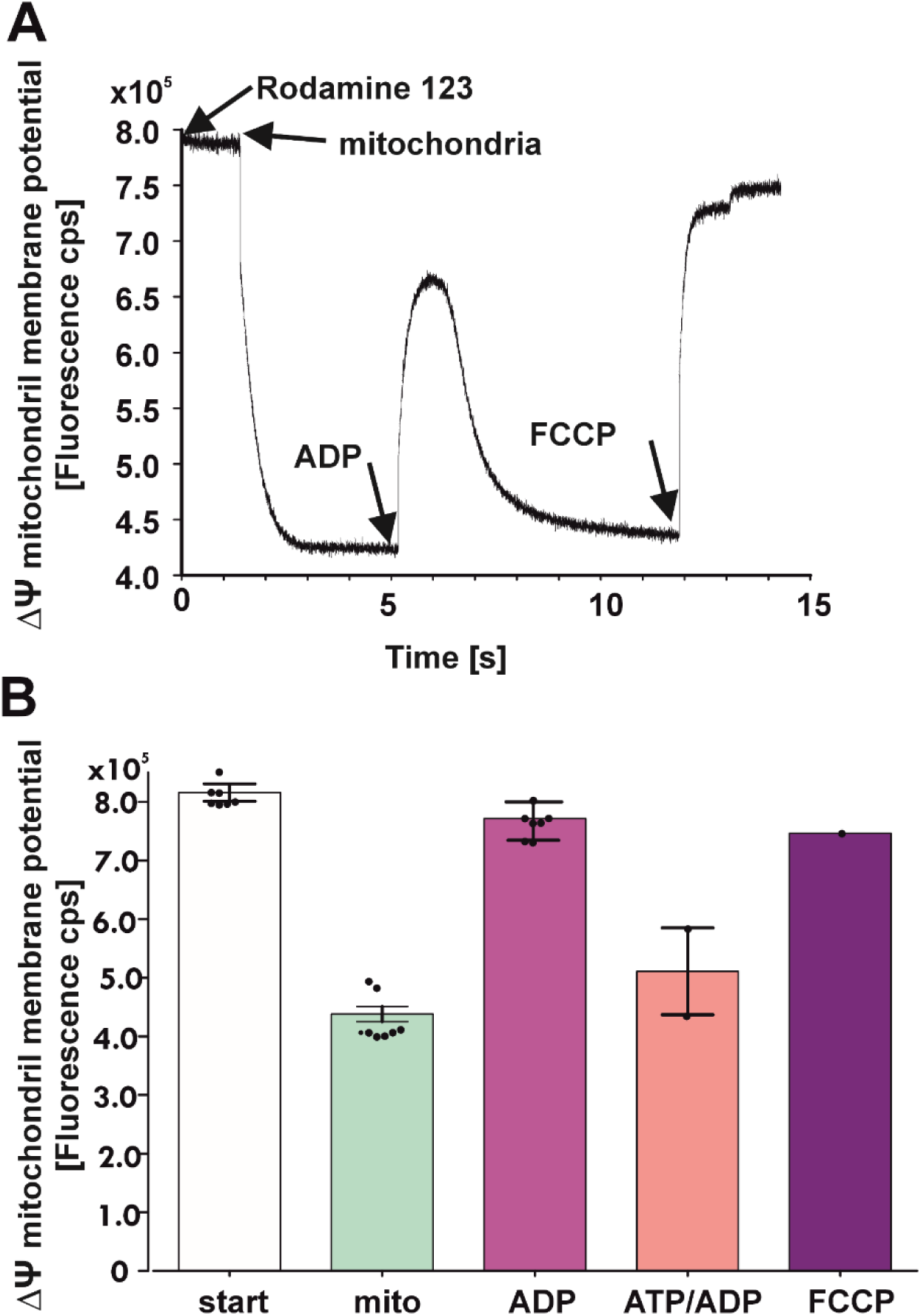
Assessment of the functional integrity of isolated mitochondria by fluorometrically measurements of Rhodamine 123 fluorescence in MiR05 buffer supplemented with respiratory substrates. ADP and FCCP influence fluorescence of 1 μM Rhodamine 123. (A) Sample recording, (B) Collective calculations.

### Respiratory Activity of Isolated Mitochondria

Mitochondrial respiration was evaluated using high-resolution respirometry (HRR). In the presence of malate, glutamate, and succinate, mitochondria exhibited low basal oxygen consumption corresponding to state II respiration (Figure 5A). Addition of ADP (100 μM) resulted in a rapid increase in oxygen consumption (state III), followed by a decline upon ADP depletion and ATP formation (state IV). Subsequent addition of FCCP (1 μM) induced maximal uncoupled respiration, reflecting the maximal respiratory capacity of the mitochondrial preparation. Quantitative analysis demonstrated a 7.39 ± 1.25-fold increase in oxygen consumption following ADP stimulation (Figure 5B), indicating preserved oxidative phosphorylation capacity.

**Figure 5.**
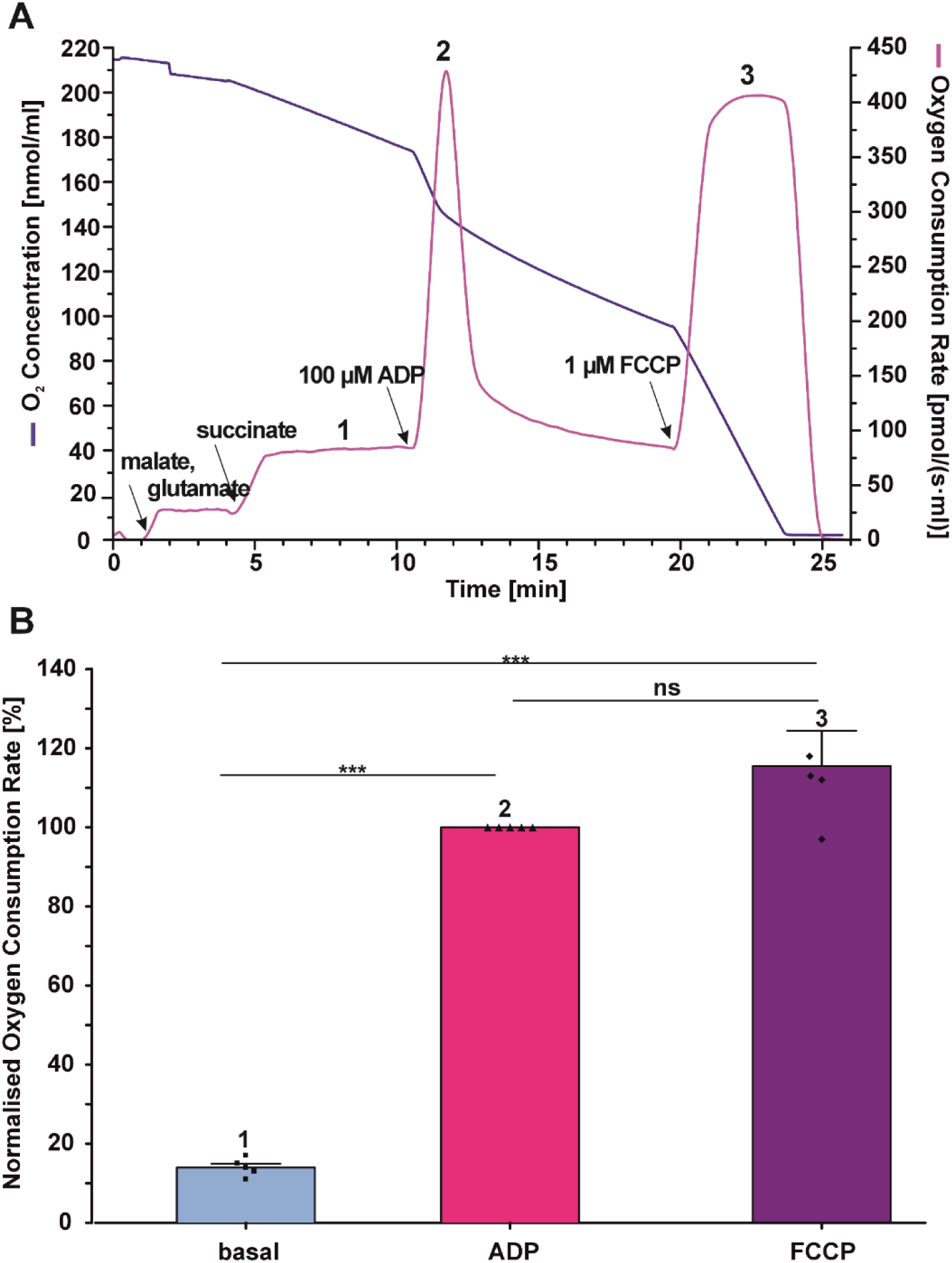
Oxygen consumption rate of mitochondria isolated from guinea pig cardiomyocytes after adding glutamate, malate, succinate, 100 µM ADP and 1 µM uncoupler - FCCP. Returning to basal respiration after ADP was spontaneous. (A) Representative example, (B) Bar chart of quantitative data for following states of respiration in cardiac mitochondria, n=8.

### Electrophysiological Properties of Isolated Mitochondria

Functionally active mitochondria exposed to a hypotonic buffer undergo osmotic shock, resulting in their transformation into mitoplasts. These structures appear as characteristic opalescent spherical bodies under light microscopy, often displaying a distinct dark region corresponding to residual fragments of the outer mitochondrial membrane. Mitoplasts prepared in this manner were used for patch-clamp recordings to assess single-channel potassium activity. A schematic representation of the experimental setup is shown in Figure 6A. A representative recording of potassium channel activity specifically a large-conductance, voltage- and Ca²⁺-activated potassium channel (mitoBK_Ca_) is presented in Figure 6B. The electrophysiological profile of the recorded currents, together with their dependence on Ca²⁺ concentration and sensitivity to the activator NS1619, is consistent with the properties of a large-conductance Ca²⁺-regulated mitoBK_Ca_ channel.

**Figure 6.**
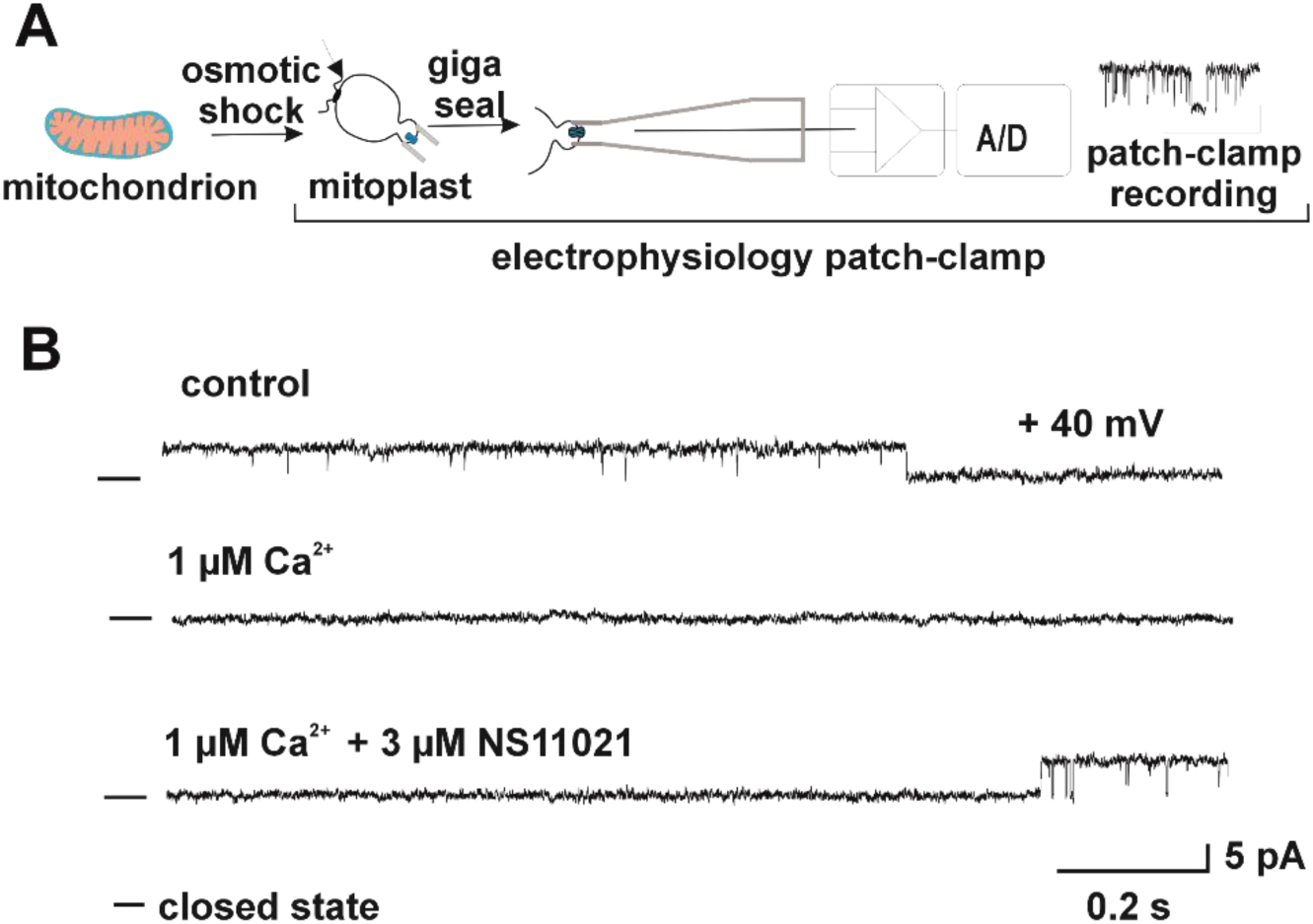
Patch-clamp of the inner membrane of mitochondria isolated from adult guinea pig cardiomyocytes. (A) Scheme of the patch-clamp recording. (B) Recording of mitoBKCa channel activity.

These findings demonstrate that mitochondria isolated using the cell strainer-based method are suitable for electrophysiological analyses. This raises the question of whether mitochondria obtained using this approach can also be applied in mitochondrial transplantation experiments in cultured cells.

### Mitochondrial Transplantation into H9c2 Cells

To determine whether mitochondria obtained using our method are suitable for transplantation into cultured cells, we employed the H9c2 cell line, derived from embryonic rat cardiac tissue and widely used in cardiovascular research. Because H9c2 cells are of rat origin, cardiomyocytes were isolated from rat hearts using a protocol analogous to that applied for guinea pig tissue and subsequently stained with the ΔΨ-sensitive fluorescent dye MitoRed. The stained cardiomyocytes exhibited a characteristic intracellular fluorescence pattern, with a periodicity of 1.55 ± 0.04 μm between intensity peaks, consistent with the typical mitochondrial organization in rat cardiomyocytes (Figure 7A). Mitochondria were then isolated from these labeled cardiomyocytes using the method developed in our laboratory and added to H9c2 cells previously stained with MitoTracker Green. Following 16 hours of incubation, the samples were analyzed using confocal microscopy (Figure 7B). H9c2 cells displayed a characteristic fluorescence pattern corresponding to mitochondrial accumulation of MitoTracker Green (Figure 7B1). After transplantation with MitoRed-labeled mitochondria, substantial internalization of exogenous mitochondria derived from rat cardiomyocytes was observed in H9c2 cells (Figure 7B2). Overlay analysis demonstrated clear colocalization of endogenous and exogenous mitochondria (green and red, respectively (Figure 7B3). That indicates successful uptake and intracellular integration of transplanted mitochondria with mitochondria of the H9c2 cells.

**Figure 7.**
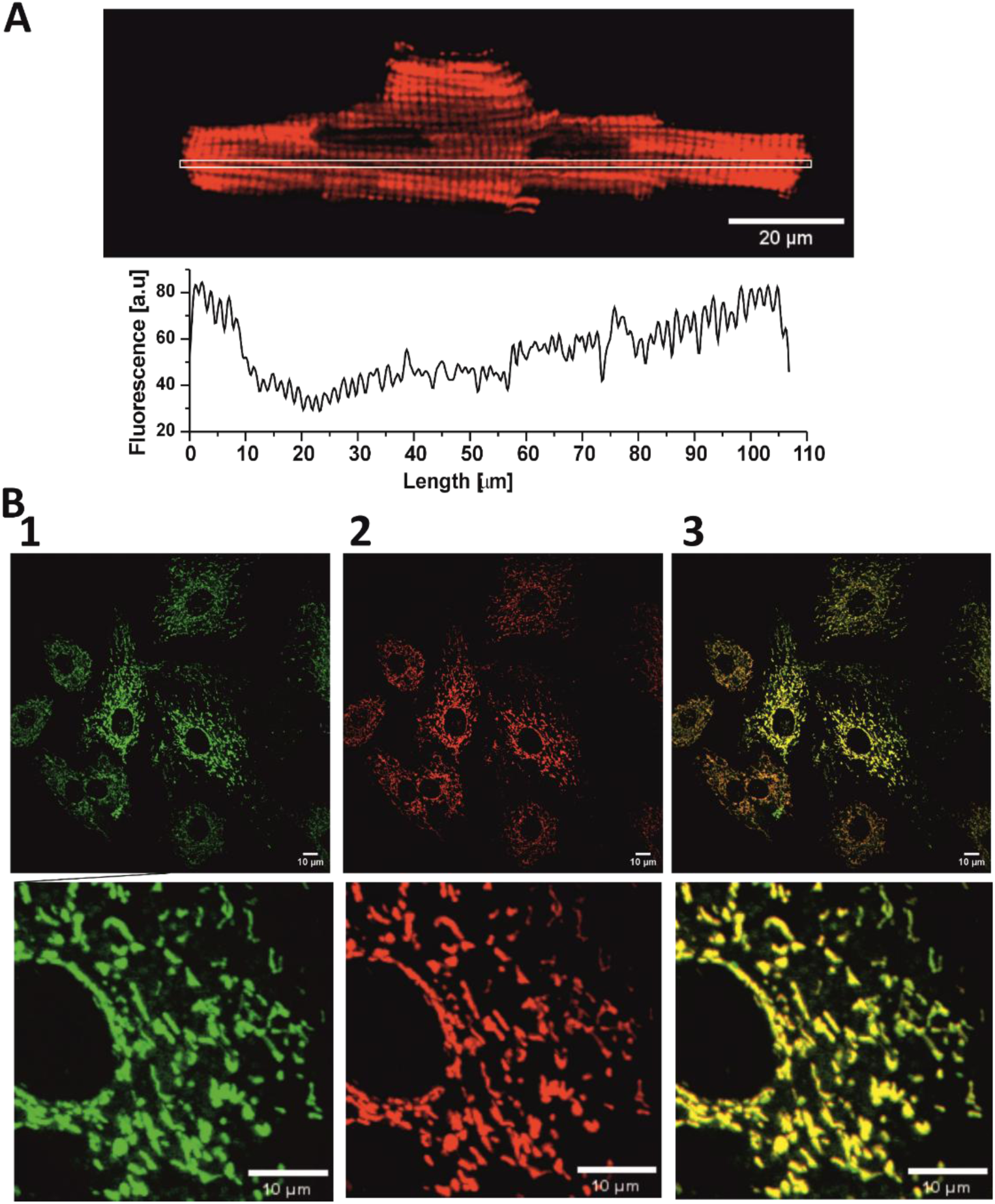
Mitochondria transfer to the H9c2 cell line. (A) Confocal image of mitochondria in rat cardiomyocyte labelled with MitoRed dye and fluorescence measurements. (B) Confocal images of H9c2 cells after mitochondrial transplantation. Mitochondria in H9c2 cells were labelled with MitoTracker Green. Rat cardiomyocytes were loaded with MitoRed dye that labelled mitochondria prior to their isolation. Images show green (1); red (2) and overlayered (3) fluorescence after rat mitochondria transplantation to H9c2 cells.

## Discussion

The manuscript presents a novel method for the isolation of mitochondria from cardiomyocytes derived from adult rodent hearts. As a studied model guinea pigs and rats were chosen. This approach employs a cell strainer to mechanically disrupt the plasma membrane of cardiomyocytes while preserving other cell types generated during enzymatic digestion of the heart in the Langendorff perfusion system (Frankenreiter et al., 2017), owing to their smaller size (Figure 1). It is important to note that cardiomyocytes constitute only approximately 25–35% of all cardiac cells; the remaining population consists predominantly of fibroblasts, as well as endothelial cells, vascular smooth muscle cells, and immune cells (Pinto, Ilinykh et al., 2016).

The need to develop a new method for mitochondrial isolation from cardiomyocytes arose from the insufficient yield obtained using conventional glass homogenizers such as Dounce or Potter-Elvehjem devices (Goldberg, 2021), as well as the risk of isolating only specific mitochondrial subpopulations, namely subsarcolemmal and interfibrillar mitochondria (Hollander, Thapa et al., 2014). An additional limitation of existing methods is the potential contamination of mitochondrial preparations by organelles originating from non-cardiomyocyte cell types, which may lead to misinterpretation of results. It is particularly important in single-channel patch-clamp experiments and biochemical assays due to differences in various protein distribution in mitochondria from distinct tissues (Szabo & Szewczyk, 2023, Szewczyk, Bednarczyk et al., 2025).

The proposed method addresses this issue at the initial stage by exploiting cell size differences to selectively eliminate non-cardiomyocyte populations. Another critical aspect of mitochondrial isolation is minimizing structural and functional damage during homogenization of both tissue and cellular preparations (Rahman, Xiao et al., 2018). The use of Dounce or Potter-Elvehjem homogenizers involves substantial shear forces and the generation of variable pressure gradients during piston movement, which depend strongly on operator technique. Mechanical tissue disruption may also induce cavitation, significantly compromising mitochondrial integrity (Gross, Greenberg et al., 2011). Similarly, digitonin-based permeabilization, although widely used for selective plasma membrane solubilization, may damage the outer mitochondrial membrane (OMM) in a concentration-dependent manner (Frezza, Cipolat et al., 2007, Wieckowski et al., 2009). This effect was observed in oxygen consumption assays, where excessive digitonin leads to reduced respiration rates. An alternative strategy involves nitrogen cavitation, where cells are exposed to high nitrogen pressure (typically 400–1000 psi) followed by rapid decompression, resulting in gas bubble formation, plasma membrane rupture, and release of mitochondria and other organelles into the buffer (Kristián, Hopkins et al., 2006, Younis, Lavery et al., 2023). Although this method provides more uniform disruption compared to mechanical homogenization, it may reduce oxygen availability and potentially inactivate pressure-sensitive enzymes. In contrast, the method developed in our laboratory for isolating mitochondria from adult rodent cardiomyocytes utilizes cell strainers with pore sizes of 40 μm or 30 μm. Given the dimensions of cardiomyocytes (approximately 140 μm in length and ∼50 μm in width), the application of controlled mechanical force using a cell scraper allows selective disruption of the plasma membrane proportional to its mechanical properties and the contractile apparatus. Simultaneously, smaller non-cardiomyocyte cells pass through the strainer and can be subsequently removed by differential centrifugation (Figure 1). Morphological and functional analyses demonstrate that mitochondria isolated using this approach retain intact ultrastructure, as confirmed by transmission electron microscopy (Figure 2 B 3, 4), comparable to that observed in situ within cardiomyocytes (Figure 2 B 2). Functionally, these mitochondria preserve their ability to generate and modulate the inner mitochondrial membrane potential (ΔΨ IMM) upon addition of respiratory substrates (Figure 4 A, B). The introduction of ADP (state 3 respiration) induces an increase in Rhodamine 123 fluorescence, reflecting ΔΨ IMM depolarization, whereas ADP phosphorylation to ATP (state 4) results in fluorescence quenching, indicating restoration of membrane potential and full mitochondrial functionality. This functional stability is maintained over several hours. The addition of the uncoupler FCCP (1 μM) induces rapid depolarization and increased fluorescence (Figure 4 A, B).

Changes in ΔΨ IMM are expected to correlate with electron transport chain activity and oxygen consumption. High-resolution respirometry revealed a substantial (>7-fold) increase in basal respiratory activity upon ADP addition in mitochondria isolated using the cell strainer method in the presence of respiratory substrates (Figure 5 A, B). Furthermore, the absence of changes in respiration following cytochrome c addition indicates preserved OMM integrity. The high purity and functional integrity of the isolated mitochondria make them particularly suitable for single-channel electrophysiological recordings using the patch-clamp technique (Figure 6 A, B). Numerous recent studies have focused on mitochondrial transplantation as a strategy to improve cellular function, particularly following ischemia/reperfusion (I/R) injury (Singh, Tahavvori et al., 2026). Mitochondria isolated from rat hearts using our method were successfully transplanted into H9c2 cells (Figure 7 B 1–3). Contrary excessively aggressive isolation procedures may disrupt mitochondrial structure and promote their elimination via mitophagy (Wang, Sun et al., 2026). Indeed, previous studies have shown that the bioenergetic benefits of mitochondrial transplantation may be transient (Ali Pour et al., 2020).

Confocal microscopy analysis demonstrated efficient uptake of Mito Red labeled mitochondria by recipient cells. This high transplantation efficiency likely reflects the preserved structural integrity and reduced proteotoxic stress associated with the isolated mitochondria (Ali Pour et al., 2021). Based on these promising results, further studies are planned to evaluate the bioenergetic improvement of recipient cells.

### Final remarks

Conventional methods for mitochondrial isolation from cardiomyocytes, including mechanical homogenization (Dounce, Potter-Elvehjem) and digitonin treatment, are limited by low yield, risk of mitochondrial damage, and contamination with mitochondria derived from non-cardiomyocyte cell types (fibroblasts, endothelial cells, smooth muscle cells, immune cells). This issue is particularly relevant given that cardiomyocytes account for only 25–35% of cardiac cells, increasing the likelihood of obtaining heterogeneous mitochondrial populations that may confound patch-clamp and biochemical analyses. The proposed method, based on the use of cell strainers (30–40 μm pore size), enables selective isolation of mitochondria almost exclusively from cardiomyocytes while smaller cells are centrifuged in their entirety, thereby enhancing the purity of mitochondrial preparations. Compared to classical approaches such as glass homogenization, digitonin permeabilization, or nitrogen cavitation, this method minimizes mechanical stress and reduces the risk of compromising mitochondrial membrane integrity. Mitochondria isolated using this technique retain full structural and functional integrity and remain competent for transplantation, which is particularly relevant in the context of therapies aimed at restoring cellular bioenergetics following ischemia/reperfusion injury.

The presented method for isolating mitochondria from cardiomyocytes is characterized by high efficiency, the ability to remove small cells whose mitochondria would otherwise constitute unwanted contaminants, minimizes mechanical damage, and preserves mitochondrial functionality. The resulting mitochondria can be used for both bioenergetics and transplantation research, overcoming the limitations of existing isolation techniques.

## Authors’ contributions

Participated in research design: Lewandowska, Kalenik, Szewczyk, Wrzosek

Conducted experiments: Lewandowska, Kalenik, Wrzosek

Wrote or contributed to the writing of the manuscript: Lewandowska, Kalenik, Szewczyk, Wrzosek

## Acknowledgements

The study was supported by the Polish National Science Centre (MAESTRO grant No. 2019/34/A/NZ1/00352 to A.S.). TEM studies were performed in the Laboratory of Electron Microscopy of the Nencki Institute, supported by the project financed by the Minister of Science and Higher Education based on contract No 2022/WK/05 (Polish Euro-BioImaging Node “Advanced Light Microscopy Node Poland”).

## Conflict of interest statement

The authors declare no conflicts of interest.

## Notes

https://infraredmito.nencki.edu.pl/

### Competing Interest Statement

The authors have declared no competing interest.

